# Small-scale spatial structure affects predator-prey dynamics and coexistence

**DOI:** 10.1101/2019.12.17.880104

**Authors:** Anudeep Surendran, Michael Plank, Matthew Simpson

## Abstract

Small-scale spatial variability can affect community dynamics in many ecological and biological processes, such as predator-prey dynamics and immune responses. Spatial variability includes short-range neighbour-dependent interactions and small-scale spatial structure, such as *clustering* where individuals aggregate together, and *segregation* where individuals are spaced apart from one another. Yet, a large class of mathematical models aimed at representing these processes ignores these factors by making a classical mean-field approximation, where interactions between individuals are assumed to occur in proportion to their average density. Such mean-field approximations amount to ignoring spatial structure. In this work, we consider an individual based model of a two-species community that is composed of *consumers* and *resources*. The model describes migration, predation, competition and dispersal of offspring, and explicitly gives rise to varying degrees of spatial structure. We compare simulation results from the individual based model with the solution of a classical mean-field approximation, and this comparison provides insight into how spatial structure can drive the system away from mean-field dynamics. Our analysis reveals that mechanisms leading to intraspecific clustering and interspecific segregation, such as short-range predation and short-range dispersal, tend to increase the size of the resource species relative to the mean-field prediction. We show that under certain parameter regimes these mechanisms lead to the extinction of consumers whereas the classical mean-field model predicts the coexistence of both species.

## 1 Introduction

Mathematical modelling of ecology and cell biology processes, such as predator-prey dynamics and immune cell-pathogen interactions, can provide insight into the impact of various interaction mechanisms that influence community dynamics. Traditional mathematical models are based on making a mean-field approximation, where the community is assumed to be locally well mixed and the presence of an individual at one location is independent of the presence or absence of an individual at any other location (Law and Dieckmann 2000; Law et al. 2003; Baker and Simpson 2010; Grunbaum 2012). Mean-field models, such as the Lotka-Volterra model of predator-prey dynamics and the logistic growth model do not incorporate any information about the spatial correlation between the locations of individuals. These classical mean-field models are typically written as ordinary differential equations that govern the time evolution of the average density of individuals (Edelstein-Keshet 2005). Some prey-predator models incorporate different types of functional responses, such as density dependence (Abrams and Ginzburg 2000; Wang et al. 2009), but do not explicitly consider the role of spatial structure. Spatially explicit mean-field models are also commonly used to help understand various ecology and cell biology processes, where the density dynamics is often described using partial differential equations (Murray 1989; Hillen and Painter 2009). Other types of spatially explicit models include coupled map lattices, integro-difference equations, and various kinds of metapopulation models (Britton 2003). Even though these spatially explicit models express the density of individuals as a function of spatial location, interactions between individuals are implicitly assumed to be given by local mean-field conditions described by the average density alone.

In many situations, short-range interactions are thought to play a significant role in determining community dynamics (Penczykowski et al. 2016; Galetti et al. 2018). For example, in a prey-predator community, an individual member of the prey population located nearby a cluster of predators could experience increased mortality due to the high risk of predation. Similarly, localised intraspecies competition for shared food resources could lead to an increase in the death rates of conspecific individuals. Neighbour-dependent interactions are known to lead to spatial structures (Tobin and Bjornstad 2003; Santora et al. 2010). Typical forms of spatial structures relevant to ecological and biological systems include: (i) *clustering* where individuals aggregate together; and (ii) *segregation* where individuals tend to spread out as much as possible (Binny et al. 2015; Treloar et al. 2015; Surendran et al. 2019). These types of spatial structures can occur within communities composed of single or multiple species (Markham et al. 2013; Gerum et al. 2018; Dini et al. 2018).

While spatial structure is known to impact the macroscale density dynamics of a community, commonly-used mean-field models neglect these effects. In contrast, individual based models (IBM) are useful to investigate the effects of short-scale interactions and the formation of spatial structure. Many IBMs are lattice-based, where the interactions between individuals are dictated by an artificial lattice structure (Mobilia et al. 2006; Mobilia et al. 2007; Baker and Simpson 2010; Dobramysl and Tauber 2013). A more realistic approach is to consider lattice-free IBMs, where interactions can be incorporated more realistically in a spatially continuous framework. The IBM considered in this study is an extension of our previous work (Surendran et al. 2018) examining the neighbour-dependent interactions between motile agents (e.g. cells) and stationary agents (e.g. obstacles) in the context of experimental cell biology. Here, we extend this framework to accommodate interactions relevant to ecological processes, such as predation and competition. Similar IBMs have been used previously to study the impact of motility and fecundity rates on density dynamics (Murrell 2005) as well as exploring the impact of spatial patterns on the evolution and natural selection of prey dispersal (Barraquand and Murrell 2012).

In this work, we explore various spatial structure forming mechanisms such as short-range predation, short-range-dispersal of offspring and short-range intraspecies competition in a community of consumers and resources. We examine the effects of these mechanisms, both in isolation and in combination, and specifically examine how these effects generate spatial structures that lead to deviations from mean-field dynamics. These investigations are conducted by comparing IBM simulations with the solution of appropriate mean-field equations. The spatial configuration of the community is analysed in terms of pair correlation functions (PCF) (Agnew 2014; Binny et al. 2016a, b). The PCFs measure spatial structure within and between species. This analysis assists our interpretation of the impact of spatial structure. We demonstrate scenarios where the mean-field model completely fails to capture the spatial effects and predict qualitatively different behaviours.

## 2 Individual-based model

We consider a community that consists of two distinct species, namely consumers and resources. The consumers are a group of individuals undergoing proliferation, death and movement events. They can be thought of as ecological predators or immune cells (Abrams 2000; Akira et al. 2006; Soehnlein et al. 2017). These consumers predate on the resources. The resources are also motile and proliferative. The resources can be thought of as a population of ecological prey or biological pathogens (Rincon et al. 2017; Hunt and Brown 2018; Vijay 2018). In our model, consumers and resources are distributed on a continuous two dimensional domain of size *L* × *L* with population sizes *N*_*c*_(*t*) and *N*_*r*_(*t*), respectively. The IBM is constructed for spatially homogeneous problems, where the population density in a small region, averaged over multiple realisations of the IBM, is independent of the location of that region (Plank and Law 2015). This means our framework is relevant to communities that do not have macroscopic gradients in the density of individuals (Jin et al. 2018).

The death rate of *n*^th^ resource individual located at 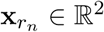, where *n* = 1, 2, …, *N*_*r*_(*t*), is taken to be a result of both interspecies predation and intraspecies competition that arise from interactions with other individuals in the population,

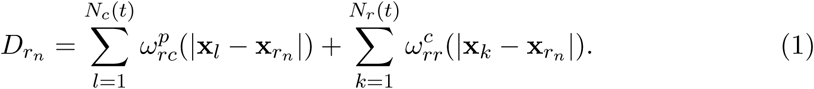

The first term on the right of Equation (1) is the death rate from predation by consumers, specified by the kernel 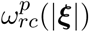. The second term on the right of Equation (1) accounts for the contribution of competition to the resources death rate, specified by the kernel 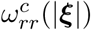. Both kernels in Equation (1) are decreasing functions of distance, |***ξ***|. A schematic representation summarising the impact of predation and competition on the event rates of individuals is given in Figure 1.

**Figure 1:**
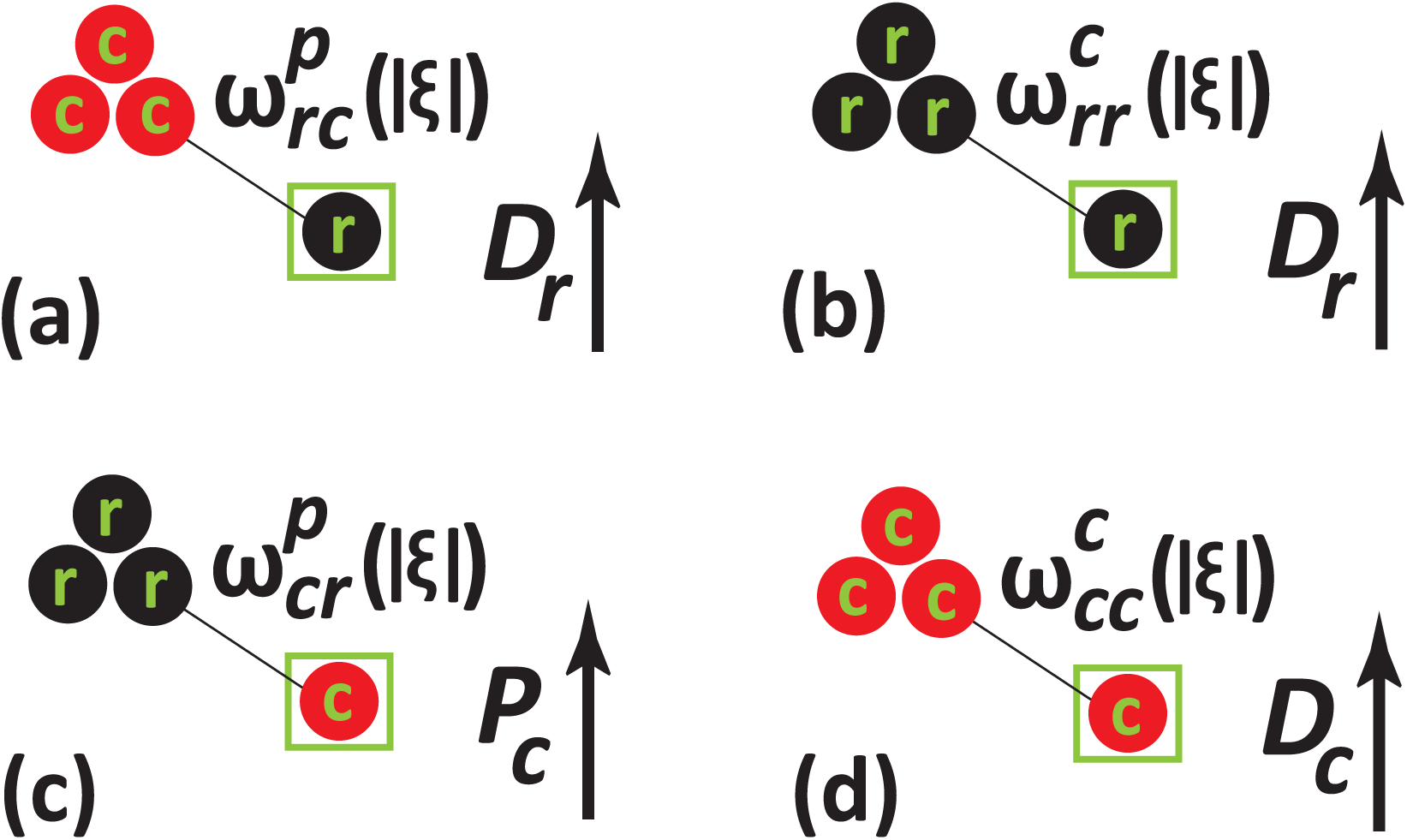
Schematic representation of how neighbour-dependent interactions affect proliferation and death rates of resources (black) and consumers (red). The green squares indicate the reference individual whose event rates are altered by the interactions. **a** impact of predation interaction on a resource leading to an increased death rate, *D*_*r*_. **b** impact of intraspecies competition of resources, leading to an increased death rate, *D*_*r*_. **c** impact of predation interaction on consumers, leading to an enhanced the proliferation rate, *P*_*c*_. **d** impact of intraspecies competition of consumers, leading to an increase in the death rate, *D*_*c*_.

We consider Gaussian interaction kernels to ensure the strength of interaction between individuals decreases with separation distance. The predation kernel describing the contribution of a neighbouring consumer at a displacement, ***ξ***, to the death rate of a reference resource individual is given by,

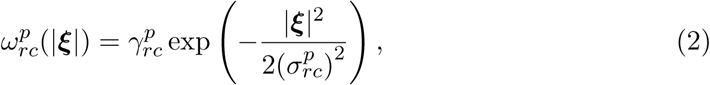

where 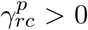 and 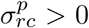 represent the predation strength and the spatial extent of predation, respectively. There is no strict requirement on the functional form of the interaction kernel. Non-Gaussian kernels can be incorporated into the IBM, depending on the specific applications (Plank et al. 2019). Competition kernels also take the same functional form. For simplicity, we assume that resources proliferate at a constant rate, 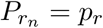, and that this rate is unaffected by the presence of other individuals.

Consumer reproduction is taken to be dependent on the predation and subsequent consumption of resources, which provide the required biomass for offspring. We use a predation kernel, 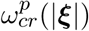, to account for the contribution of resources in the enhancement of the proliferation rate of consumers. Setting 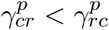 reflects a realistic situation where the consumption of a single resource is insufficient to generate a single consumer. We also make a reasonable assumption that, 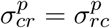, so that the distance over which consumers and resources interact is symmetric. The proliferation rate of consumers is computed by summing the contributions from all the resources,

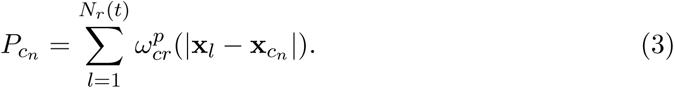

The mortality of consumers depends on competitive interactions between consumers. The death rate of consumers is given by,

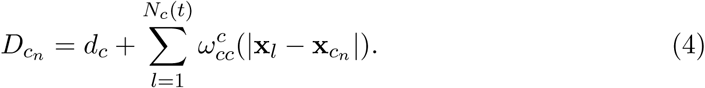

The first term on the right of Equation (4) is a constant intrinsic death rate of consumers which is unaffected by the presence of other individuals in the model. The second term on the right of Equation (4) accounts for the contribution of competition between consumers. For simplicity we assume the movement rate of consumers and resources are constant, and given by *m*_*c*_, and *m*_*r*_, respectively.

The IBM is simulated using the Gillespie algorithm (Gillespie 1977). We use periodic boundary conditions over the computational domain since the domain considered is large enough to avoid any edge effects. Initial population sizes of consumers and resources are *N*_*c*_(0) and *N*_*r*_(0), respectively, and the locations of these individuals are initially chosen at random. At each time step, the neighbour-dependent event rates of individuals are computed according to Equation (1) and Equations (3)–(4). The total event rate of all individuals is computed by summing all the intrinsic and neighbour-dependent rates as,

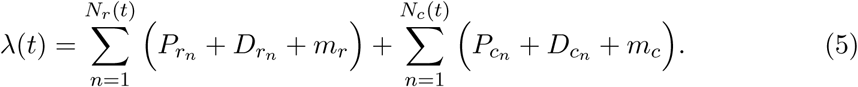

The time intervals between successive events are distributed according to an exponential distribution with mean 1*/λ*(*t*). At each time increment of the Gillespie algorithm, one of the three possible events occurs. The probability of the occurrence of an event is proportional to the rate of that event (Baker and Simpson 2010). When a consumer proliferates, a daughter consumer is placed at a displacement, ***ξ***, drawn from a bivariate normal distribution with mean zero and standard deviation, 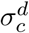, resulting in an increase of the consumer population by one. Here, 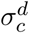 is the consumer dispersal range. Similarly, the proliferation of a resource individual involves placing a daughter resource at a displacement sampled from a bivariate normal distribution with mean zero and standard deviation 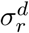. When a consumer undergoes a movement event, it travels a displacement (|***ξ***| cos(*θ*), |***ξ***| sin(*θ*)), where the distance moved by a consumer, |***ξ***|, is drawn from a truncated Gaussian distribution with a positive mean 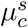 and a narrow standard deviation 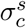, such that 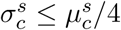. To ensure the movement distance is always positive, we truncate the Gaussian at distances greater than 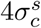 from the mean. The direction of movement, *θ* ∈ [0, 2*π*], is uniformly distributed. Movement of a resource is specified by a similar Gaussian distribution with mean 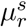 and standard deviation 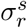.

To analyse the resulting community dynamics, we calculate the average density of consumers and resource by dividing the number of individuals in both the species with the area of the domain as, *Z*_*c*_(*t*) = *N*_*c*_(*t*)*/L*^2^ and *Z*_*r*_(*t*) = *N*_*r*_(*t*)*/L*^2^, respectively. The average density of pairs of individuals reveals the correlation between the locations of individuals (Bolker and Pacala 1999; Law et al. 2009; Ovaskainen et al. 2014). We calculate several PCFs, defined as the average densities of pairs normalised by the density of pairs in a community without any spatial structure (i.e. a community that evolves in such a way that the mean-field assumption is valid). We define the PCF as a function of the separation distance between pairs of individuals, |***ξ***|, and time, *t*. Since there are two species in the model, there are three unique PCFs: (i) the auto-PCF of consumers, *C*_*cc*_(|***ξ***|, *t*); (ii) the auto-PCF of resources, *C*_*rr*_(|***ξ***|, *t*), and, (iii) the cross-PCF, *C*_*cr*_(|***ξ***|, *t*), respectively. In the complete absence of spatial structure we have *C*_*cc*_(|***ξ***|, *t*) = *C*_*rr*_(|***ξ***|, *t*) = *C*_*cr*_(|***ξ***|, *t*) = 1. When the auto-PCF of consumers is greater than unity, *C*_*cc*_(|***ξ***|, *t*) > 1, we have a larger number of pairs of consumers separated by a distance, |***ξ***|, than we would have in a community with spatially random configuration. We refer to this configuration as clustered. When *C*_*cc*_(|***ξ***|, *t*) < 1, we have a smaller number of pairs of consumers separated by a distance, |***ξ***|, than we would have in a community with spatially random configuration and this situation corresponds to a segregated spatial structure. Similarly, clustered and segregated spatial patterns of resources correspond to *C*_*rr*_(|***ξ***|, *t*) > 1 and *C*_*rr*_(|***ξ***|, *t*) < 1, respectively. Interspecies clustering and segregation of consumers and resources correspond to *C*_*cr*_(|***ξ***|, *t*) > 1 and *C*_*cr*_(|***ξ***|, *t*) < 1, respectively.

To compute the auto-PCF of consumers, we consider one consumer individual as a reference individual and then calculate the distance between the reference individual and all other *N*_*c*_(*t*) − 1 consumers. We record the distances between the reference individual and other individuals of the same species. The auto-PCF is constructed by counting the distances that fall into the interval [|***ξ***|−Δ|***ξ***|*/*2, |***ξ***|+Δ|***ξ***|*/*2] (Binder and Simpson, 2015). The bin count is normalized by a factor of 2*π*|***ξ***|Δ|***ξ***|*N*_*c*_(*N*_*c*_ − 1)*/L*^2^ to ensure that *C*_*cc*_(|***ξ***|, *t*) = 1 in the absence of spatial structure. A similar procedure is followed to compute *C*_*rr*_(|***ξ***|, *t*) and *C*_*cr*_(|***ξ***|, *t*).

### 2.1 Mean-field dynamics

Here, we present a continuum description of the IBM by invoking the mean-field assumption. We denote the density of species *i* at time *t*, averaged over many identically-prepared realisations of the IBM as *Z*_*i*_(*t*). Note that, in our model, average density is independent of location owing to the combination of boundary conditions and initial condition used. In the IBM, the presence of other individuals in the neighbourhood of a focal individual is affected by the community’s spatial structure. However, the mean-field assumption ignores this dependence by assuming that the probability that there is an individual of species *i* at any given location ***ξ*** at time *t* is independent of the occupancy of other individuals, and is given by *Z*_*i*_(*t*). For example, the expected proliferation rate of a consumer at location ***ξ*** and time *t* is, 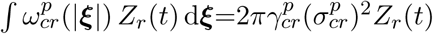. Approximating the other birth and death rates in the community in a similar way leads to

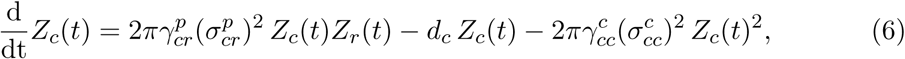

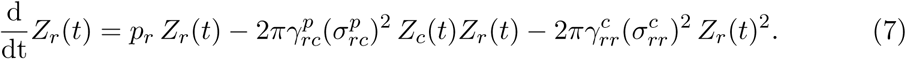

This system of equations can be derived rigorously from the IBM under the mean-field assumption by replacing the second moment of the spatial point process with the product of the first moments (Plank and Law, 2015), but we omit the details of this here. In this work we study the solution of Equations (6)–(7) numerically using the ode45 routine in MATLAB (Mathworks 2019).

## 3 Results

In this section, we compare averaged data from IBM simulations with the solution of the mean-field model to explore the impact of spatial structure on the community dynamics. The auto- and cross-PCFs help to distinguish between different spatial structures formed due to the intraspecies and interspecies interactions. Linking the PCFs and averaged densities from the IBM with the solution of the mean-field model provides comprehensive insight into the role of spatial structure in driving community dynamics. We first present simulation results that are in good agreement with the solutions of the mean-field model. We then use these initial results as a benchmark for exploring other parameter combinations. Mean-field dynamics are replicated in the IBM by considering sufficiently large predation, competition and dispersal ranges. Under these conditions the spatial configuration of individuals does not strongly influence community dynamics, and the individuals interact weakly leading to negligible correlations. A summary of parameter values for this first case are given in Table 1.

**Table 1:**
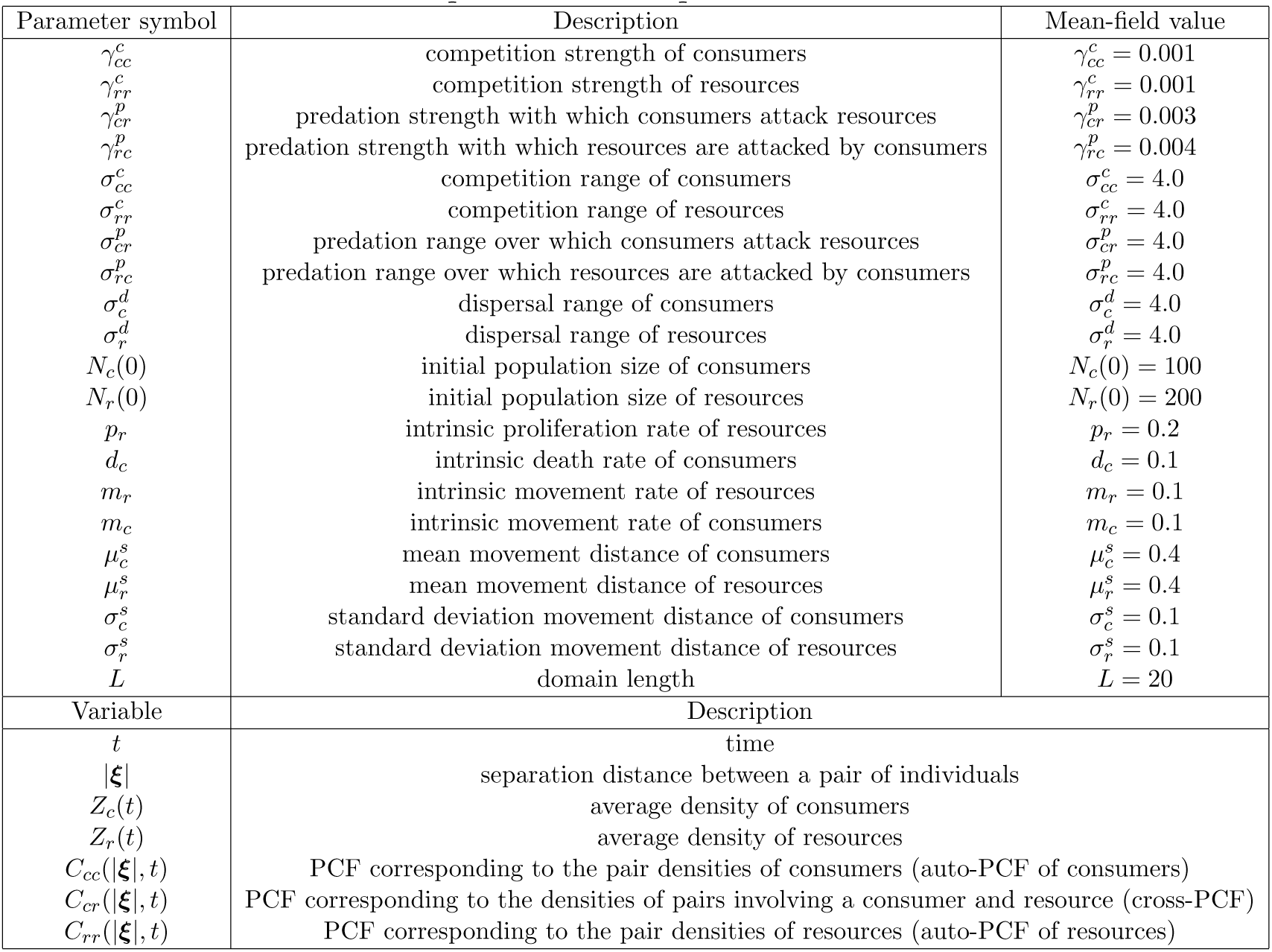
Description of model parameters and variables

In Figure 2, we show two scenarios where averaged simulation results from the IBM match well with the solution of the mean-field model. The first case, shown in Figure 2(a)–(d), corresponds to a scenario that exhibits co-existence of consumers and resources. In this case, we consider a community of consumers and resources distributed uniformly with *N*_*c*_(0) = 100 and *N*_*r*_(0) = 200, respectively, as shown in Figure 2(a). After a sufficiently long time, *t* = 200, we observe that the densities of consumers and resources and the three PCFs appear to become steady. The snapshot of the IBM at *t* = 200 in Figure 2(b) does not show any obvious spatial structure, and the auto- and cross-PCFs are approximately unity in Figure 2(d), confirming the absence of spatial structure. The temporal dynamics of the community are tracked by plotting the average densities of consumers and resources as a function of time, in Figure 2(c). Here, the mean-field model accurately matches the IBM results. These observations are expected since we consider long-range interactions and long-range dispersal.

**Figure 2:**
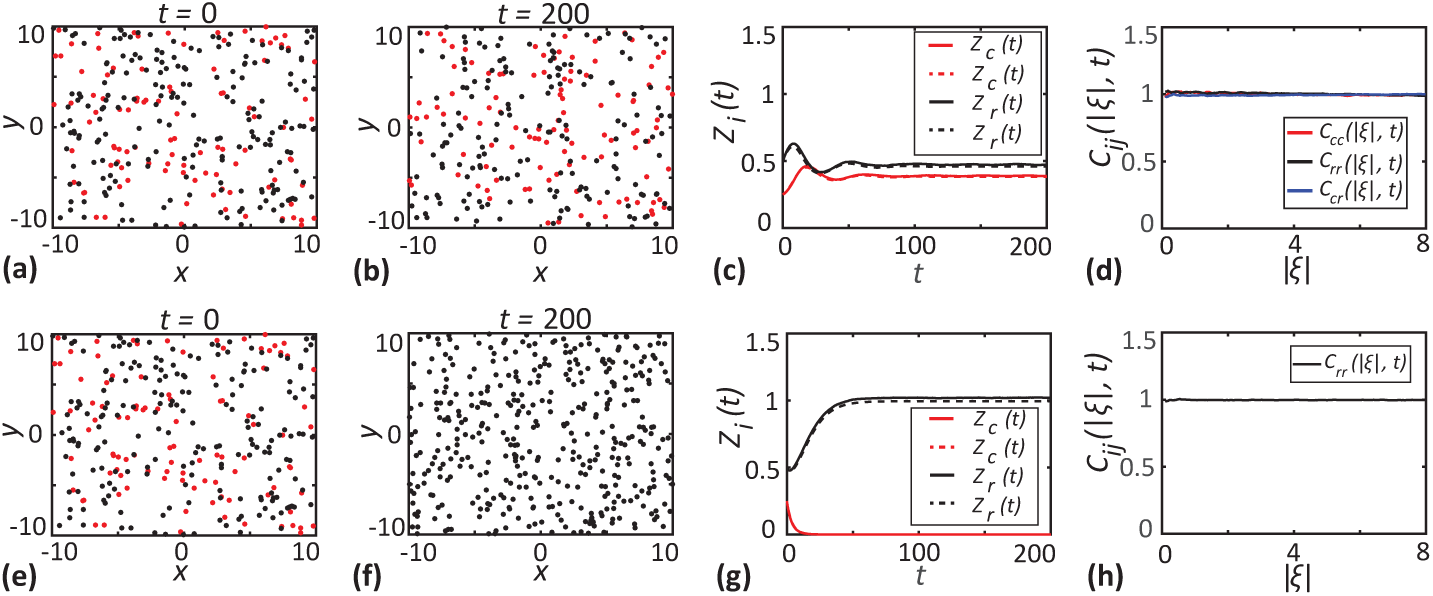
Simulation results that match well with the solutions of the mean-field model. Results in **a-d** correspond to a case where consumers and resources coexist (*p*_*r*_ = 0.2, *d*_*c*_ = 0.1). Results in **e-h** correspond to a case where consumer extinction occurs (*p*_*r*_ = 0.1, *d*_*c*_ = 0.4). **a, e** show locations of consumers (red dots) and resources (black dots) at *t* = 0. **b, f** show locations of consumers and resources at *t* = 200. **c, g** show the densities of consumers (red lines) and resources (black lines) as a function of time. Solid lines correspond to the averaged results from 1000 realisations of the IBM and dashed lines correspond to the solution of the mean-field model, Equations (6)-(7). **c, f** show the PCFs computed at *t* = 200 as a function of separation distance. Other parameter values are given in Table 1.

We present a different example of mean-field dynamics in Figure 2(e)–(h), where the consumer species eventually becomes extinct. Here, we choose an identical initial arrangement of individuals and interactions to that of the coexistence case in Figure 2(a)–(d). The only difference between the results in Figure 2(e)–(h) and Figure 2(a)–(d) is that here we specify a higher consumer death rate which leads to the extinction of consumers. The snapshot of the IBM at *t* = 200 in Figure 2(f) shows that only resources remain at this time. Consistent with this, the density of consumers in Figure 2(g) decays to zero by approximately *t* = 20. The estimate of average densities from IBM and mean-field model match well for this case. Results in Figure 2(h) indicate that *C*_*rr*_(*ξ, t*) ≈ 1, and this is consistent with the fact that the mean-field approximation is accurate. Note that we only give *C*_*rr*_(*ξ, t*) in Figure 2(h) since consumers are absent from the simulation by *t* = 200.

Given we have established that averaged data from the IBM is consistent with the solution of the mean-field model for the choice of parameters in Figure 2, we systematically vary the parameters in the IBM and explore how the resulting spatial structure affects the accuracy of the mean-field description. In all of these subsequent comparisons, we fix the mean-field equilibrium densities of consumers and resources. By fixing the equilibrium densities, we are able to compare different mechanisms and their impacts on the dynamics. Under coexistence, the equilibrium densities of consumers and resources are given by,

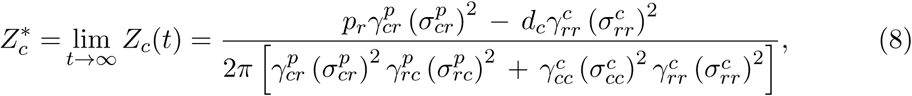

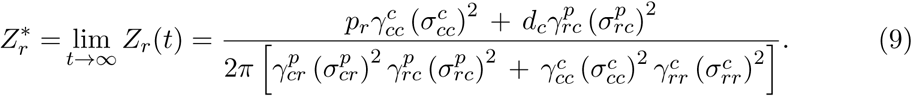

When we compare different parameter combinations, we take care to choose the interaction ranges and interaction strengths so that 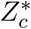 and 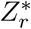 remain constant. This ensures that the long-time solution of the mean-field model is constant between the various conditions that we examine. The specific values of interaction range and strength parameters used in each of our simulations are given in respective figure captions.

### 3.1 Short range dispersal creates intraspecies clustering

Here, we investigate the impact of short-range dispersal of consumers and resources on the spatial structure of the community and how it influences the accuracy of the mean-field model. In these simulations, the predation and competition interactions are long-range, as in Figure 2. We first consider short-range dispersal of consumers, as shown in Figure 3(a)–(d). This leads to the development of intraspecies clustering among consumers, as shown in Figure 3(b), quantified by the auto-PCF with *C*_*cc*_(|***ξ***|, *t*) > 1 for small |***ξ***| in Figure 3(d). In addition we have, *C*_*rr*_(|***ξ***|, *t*) ≈ *C*_*cr*_(|***ξ***|, *t*) ≈ 1, suggesting there is no intraspecies spatial structure among resources and no interspecies spatial structure. Since the dispersal of resources is long-range, there is little correlation between the locations of parent and daughter resources, which explains why there is no intraspecies spatial structure among resources. Even though consumers exhibit strong clustering we do not see a large discrepancy between the IBM simulations and the solution of the mean-field model. This observation is due to the long-range predation and competition interactions, under which local spatial structure does not influence the dynamics of the community.

**Figure 3:**
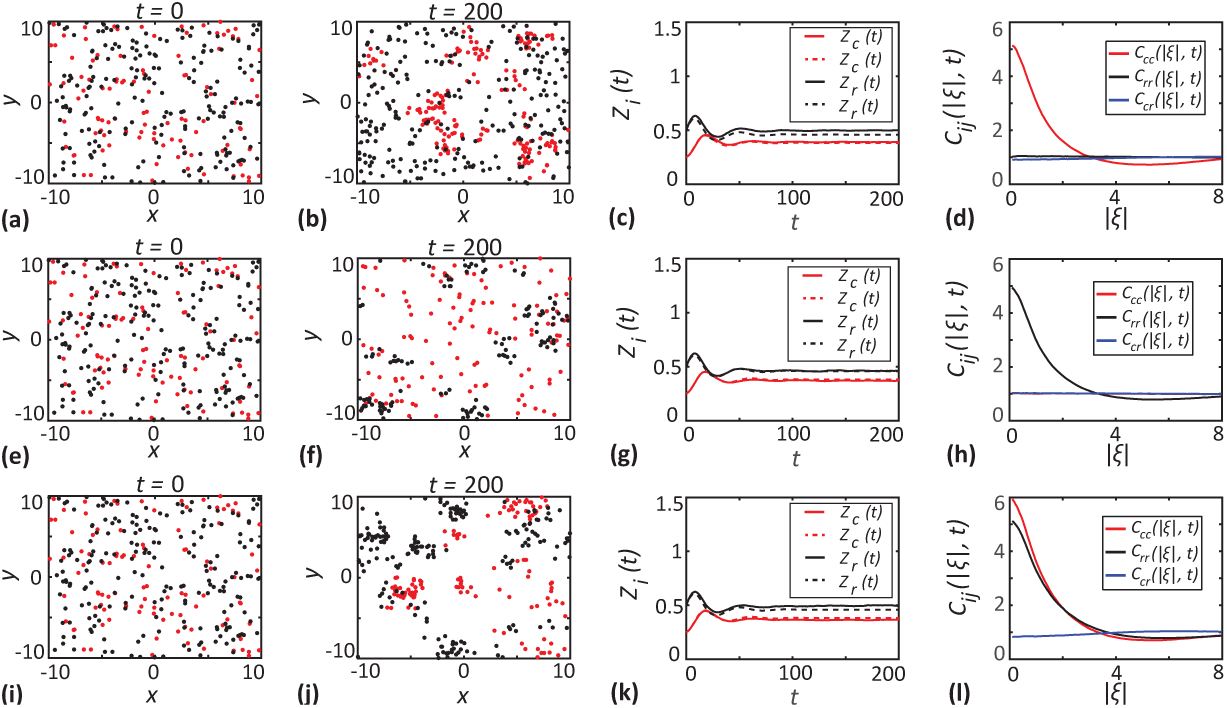
Short-range dispersal. Results in **a-d** correspond to short range dispersal of consumers 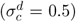. Results in **e-h** correspond to short range dispersal of resources 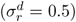. Results in **i-l** correspond to short range dispersal of both species 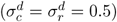. **a, e, i** show the locations of consumers (red dots) and resources (black dots) at *t* = 0. **b, f, j** show the locations of consumers and resources at *t* = 200. **c, g, k** show the densities of consumers (red lines) and resources (black lines) as a function of time. Solid lines correspond to the averaged results from 1000 realisations of the IBM and dashed lines correspond to the solution of the mean-field model, Equations (6)-(7). **d, h, l** show the PCFs computed at *t* = 200 as a function of separation distance. All other parameters are same as in Table 1.

Next, we consider the short-range dispersal of resources in Figure 3(e)–(h). After a sufficiently long time, *t* = 200, we see more resource individuals at close distances compared to the initial spatial random distribution. Results in Figure 3(h) show *C*_*cc*_ > 1, confirming clustering of resources. Again, the cluster formation is due to short-range dispersal. Similar to the case of short-range dispersal of consumers, we do not observe strong interspecies spatial structure or any discrepancy between the mean-field model and IBM estimates of the densities of consumers or resources. Finally, we consider the short-range dispersal of both consumers and resources in Figure 3(i)–(l). Since resource and consumer parents now place their offspring close to them, we see the development of intraspecies clusters of both consumers and resources.

### 3.2 Short range competition creates intraspecies segregation

We first consider short-range competition among consumers in Figure 4(a)–(d). After a sufficiently long time, *t* = 200, consumers tend to stay apart from each other, forming an intraspecies segregated spatial pattern, confirmed by *C*_*cc*_(|***ξ***|, *t*) < 1 in Figure 4(d). The intense competition at short distances increases the death rates of nearby consumers, leading to a configuration where consumers are well-separated. In addition we have *C*_*rr*_(|***ξ***|, *t*) ≈ *C*_*cr*_(|***ξ***|, *t*) ≈ 1, indicating there is no intraspecies spatial structure in resources and no interspecies spatial structure among consumers and resources. Since we consider long-range predation for this case, the consumer-resource interspecies interactions are similar to that of the mean-field case in Figure 2. Hence, the spatial structure of consumers does not impact density dynamics, as evident from the good agreement between the IBM results and the solution of the mean-field model in Figure 4(b).

**Figure 4:**
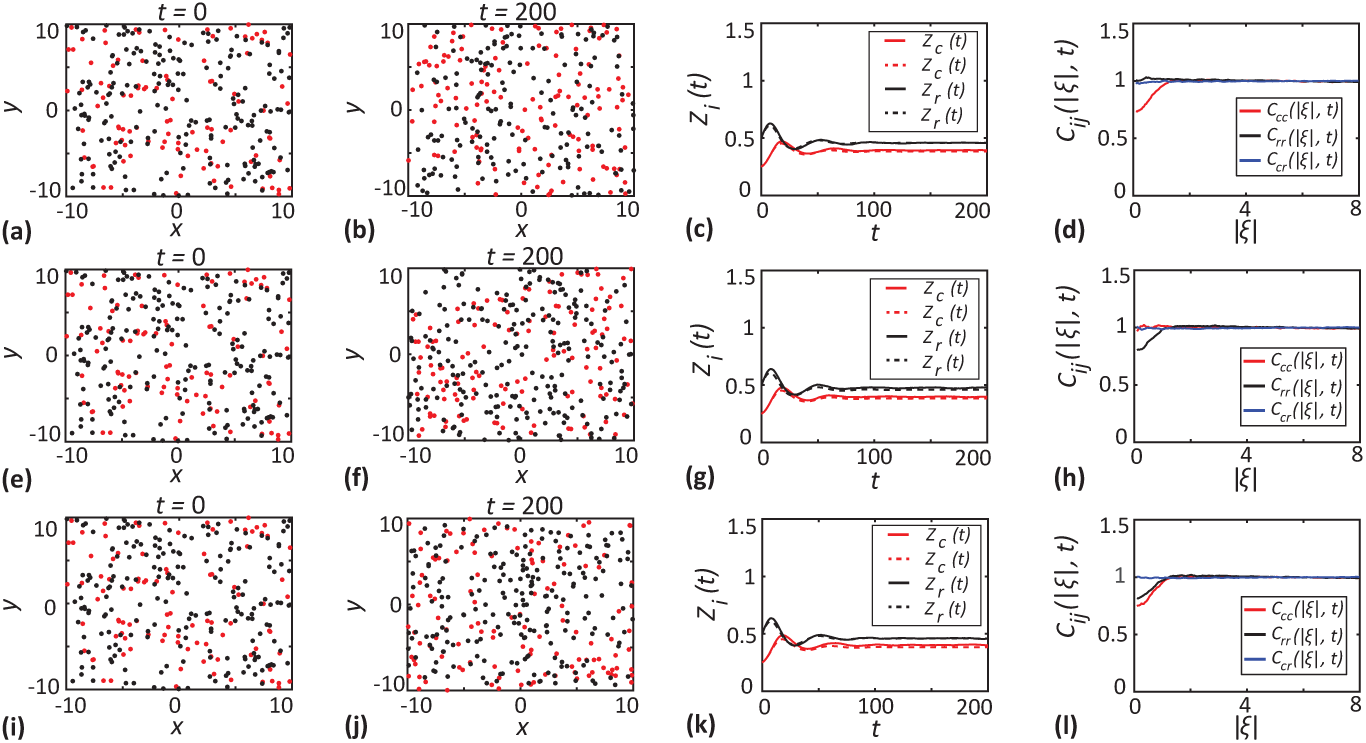
Short-range competition. Results in **a-d** correspond to short-range competition of consumers (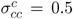 and 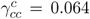). Results in **e-h** correspond to short-range competition of resources (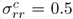 and 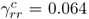). Results in **i-l** correspond to short-range competition of both species (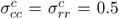 and 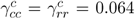). **a, e, i** show the locations of consumers (red dots) and resources (black dots) at *t* = 0. **b, f, j** show the locations of consumers and resources at *t* = 200. **c, g, k** show the densities of consumers (red lines) and resources (black lines) as a function of time. Solid lines correspond to the averaged results from 1000 realisations of the IBM and dashed lines correspond to the solution of the mean-field model, Equations (6)-(7). **d, h, l** show the PCFs computed at *t* = 200 as a function of separation distance. All other parameters are same as in Table 1.

Results in Figure 4(e)-(h) and Figure 4(i)-(l) consider intraspecies competition of resources and intraspecies competition of both species. In both cases, we observe competition-induced segregation. In Figure 4(h), *C*_*cc*_(|***ξ***|, *t*) < 1, confirms the segregation of resources, whereas *C*_*cc*_(|***ξ***|, *t*) < 1 and *C*_*rr*_(|***ξ***|, *t*) < 1, respectively in Figure 4(l) confirm the interspecies segregation between consumers and resources. Again, we observe that the density dynamics from IBM is in good agreement with the solution of the mean-field model.

### 3.3 Short range predation drives interspecies segregation

In this section we explore the impact of short-range predation. In these simulations we fix the range of competition and dispersal to be the same as in Figure 2 so that any spatial structure is driven by the influence of short-range predation. Results in Figure 5(a)-(b) show the initial spatially random configuration of individuals and the long-time outcome of the IBM, respectively. In this case, visually identifying the nature of the long-time steady spatial structure of the community is difficult. The PCFs shown in Figure 5(d), provide insight into the spatial structure. Since the predation is short-range, there is a high probability that each consumer predates upon nearby resources. Hence, the short-range predation by a consumer eventually depletes the resources locally, creating an interspecies segregation. This is confirmed by *C*_*cr*_(|***ξ***|, *t*) < 1 for short distances in Figure 5(d). There is some clustering among resources indicated by *C*_*rr*_(|***ξ***|, *t*) > 1 at small distances. This is because consumers tend to predate upon nearby resources. Those resources that are not consumed are more likely to be in regions with lower consumer density, which therefore tend to contain small clusters of resources. These spatial structures have an impact on the density dynamics. Results in Figure 5(c) show that the mean-field model underestimates the density of resources. The spatial segregation between consumers and resources assists the resources to avoid predation. Results in Figure 5(c) also show that the mean-field model underestimates the density of consumers. This may be counter-intuitive but can be attributed to the net increase in the resource population size more than offsetting the decrease in per capita contact rate between consumers and resource due to interspecific segregation.

**Figure 5:**
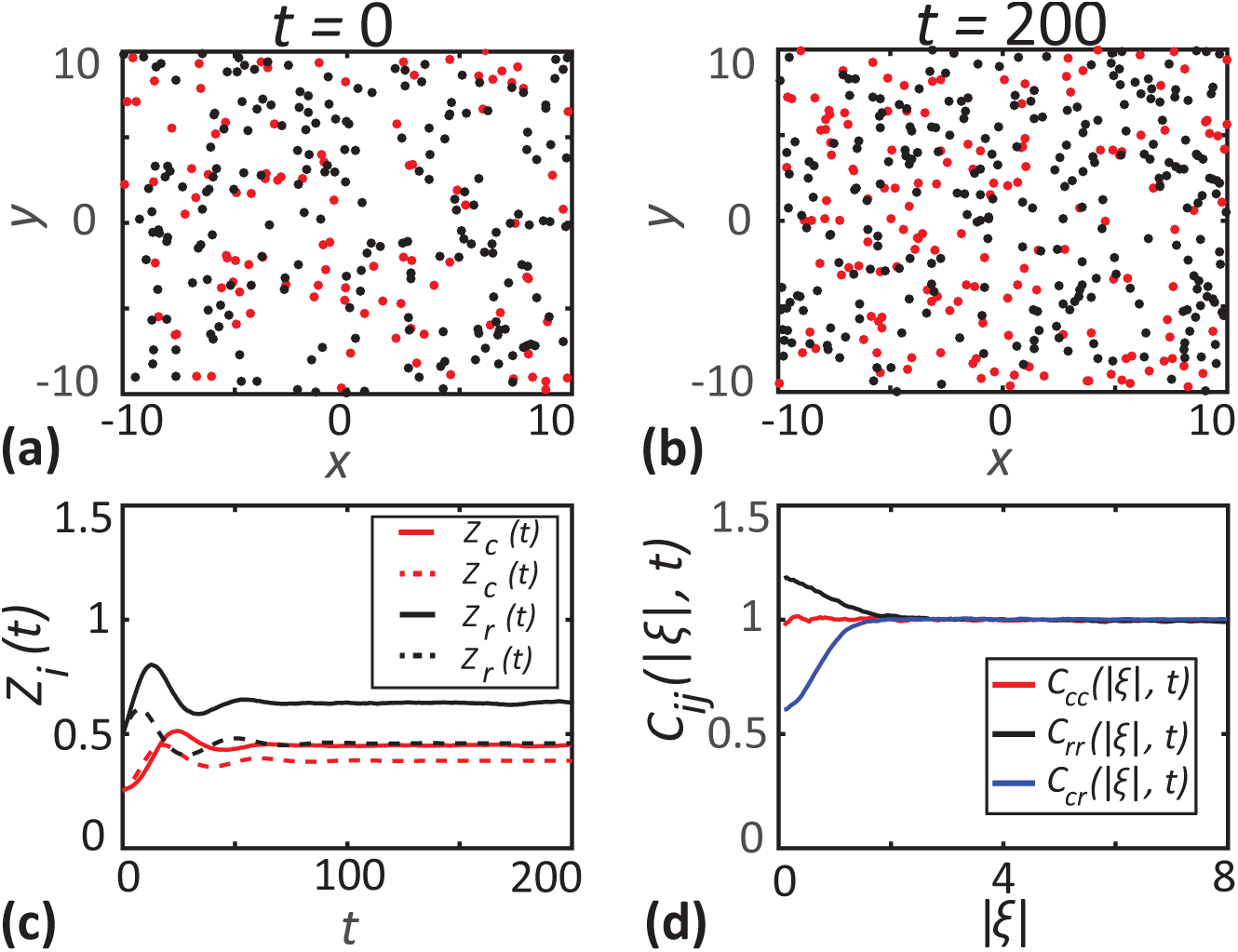
Short-range predation. **a-b** show the locations of consumers (red dots) and resources (black dots) at *t* = 0 and *t* = 200, respectively. **c** shows the densities of consumers (red lines) and resources (black lines) as a function of time. Solid lines correspond to the averaged results from 1000 realisations of the IBM and dashed lines correspond to the solution of the mean-field model, Equations (6)-(7). **d** shows the PCFs computed at *t* = 200 as a function of separation distance. Parameter values are 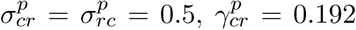 and 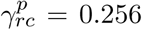. All other parameters are same as in Table 1.

### 3.4 Short-range dispersal and short-range predation enhances resources’ survival

Results in Sections 3.1–3.3 consider the impact of various mechanisms acting in isolation whereas here we investigate multiple mechanisms acting in unison. Note that, when we consider different mechanisms that generate contrasting effects on the density dynamics and spatial structure in unison, the net effect observed in the community depends on the relative strengths of those mechanisms. Hence we are always careful here to choose and report appropriate parameter values relative to previous choices in Figures 2–5, for which the opposing mechanisms have comparable strengths, such that neither is dominant over the other. If one mechanism is much stronger than others, the results will resemble the case with only that mechanism considered. We start by considering the combined effect of short-range dispersal of consumers and short-range predation in Figure 6(a)-(d). In the long-time limit we observe dispersal-induced clustering of consumers and predation-induced interspecies segregation between consumers and resources, leading to *C*_*cc*_(|***ξ***|, *t*) > 1 and *C*_*cr*_(|***ξ***|, *t*) < 1. Segregation between the two species minimises the chances of predation and results in an increase in the density of resources compared to the solution of the mean-field model. Note that the density of resources in Figure 6(c) is higher compared to the case in Figure 5, where there was only short-range predation. Since the dispersal of resources is long-range, there is a possibility of daughter resources being placed in local neighbourhoods of consumers. The probability of this happening is significantly lower when consumers are clustered rather than uniformly distributed all over the domain as in the case in Figure 5.

**Figure 6:**
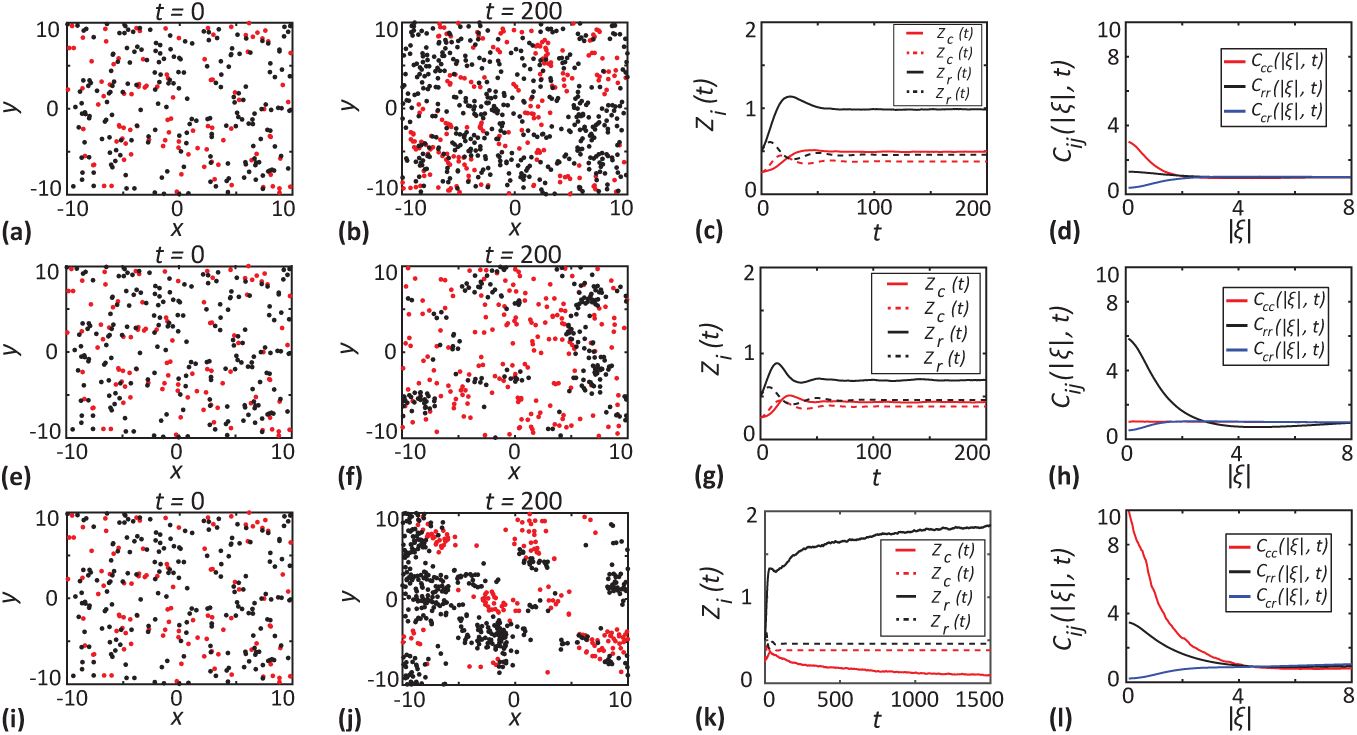
Short-range dispersal and short-range predation. Results in **a-d** correspond to short range dispersal of consumers 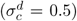 and short-range predation (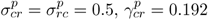 and 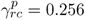). Results in **e-h** correspond to short-range dispersal of resources 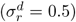 and short-range predation. Results in **i-l** correspond to short-range dispersal of both species 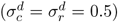 and short-range predation. **a, e, i** show the locations of consumers (red dots) and resources (black dots) at *t* = 0. **b, f, j** show the locations of consumers and resources at *t* = 200. **c, g, k** show the densities of consumers (red lines) and resources (black lines) as a function of time. Solid lines correspond to the averaged results from 1000 realisations of the IBM and dashed lines correspond to the solution of the mean-field model, Equations (6)-(7). **d, h, l** show the PCFs computed at *t* = 200 as a function of separation distance. All other parameters are same as in Table 1.

Next, we consider short-range dispersal of resources with short-range predation in Figure 6(e)–(h). These conditions give rise to dispersal induced clustering of resources and predation induced interspecies segregation, where *C*_*rr*_(|***ξ***|, *t*) > 1 and *C*_*cr*_(|***ξ***|, *t*) < 1, respectively. The density of consumers and resources from the IBM is greater than the solution from the mean-field model, but the deviation is smaller than in the previous case with short-range dispersal of consumers and short-range predation. Here the consumers are not concentrated on specific regions as compared to the previous case. Hence, the probability of predation is, on average, higher compared to the previous case.

Finally, we consider short-range dispersal of both consumers and resources along with short-range predation in Figure 6(i)–(l). In this case we see distinct clusters of consumers and resources due to short-range dispersal of both the species, confirmed by *C*_*cc*_(|***ξ***|, *t*) > 1 and *C*_*rr*_(|***ξ***|, *t*) > 1. Here, we observe an increase in the density of resources and a decrease in the density of consumers. Since both the species form clusters and occupy specific regions in the domain, the chances of pairs of consumers and resources separated by short distances is significantly reduced. The short-range predation under these circumstances leaves the consumers with reduced availability of resources for their survival. In contrast, resource grows rapidly due to the reduced risk of predation.

### 3.5 Effect of short range predation and short range competition

We consider the combined effect of short-range competition of consumers and short-range predation in Figure 7(a)–(d). Here, we see consumers tend to become isolated from other consumers and resources. While short-range predation creates interspecies segregation, the competition within the consumer subpopulation results in intraspecies segregation, confirmed by *C*_*cr*_(|***ξ***|, *t*) < 1 and *C*_*cc*_(|***ξ***|, *t*) < 1, respectively. These results indicate that the density of resources tends to be higher than the solution of the mean-field model since segregation of consumers and resources reduces the effects of predation.

**Figure 7:**
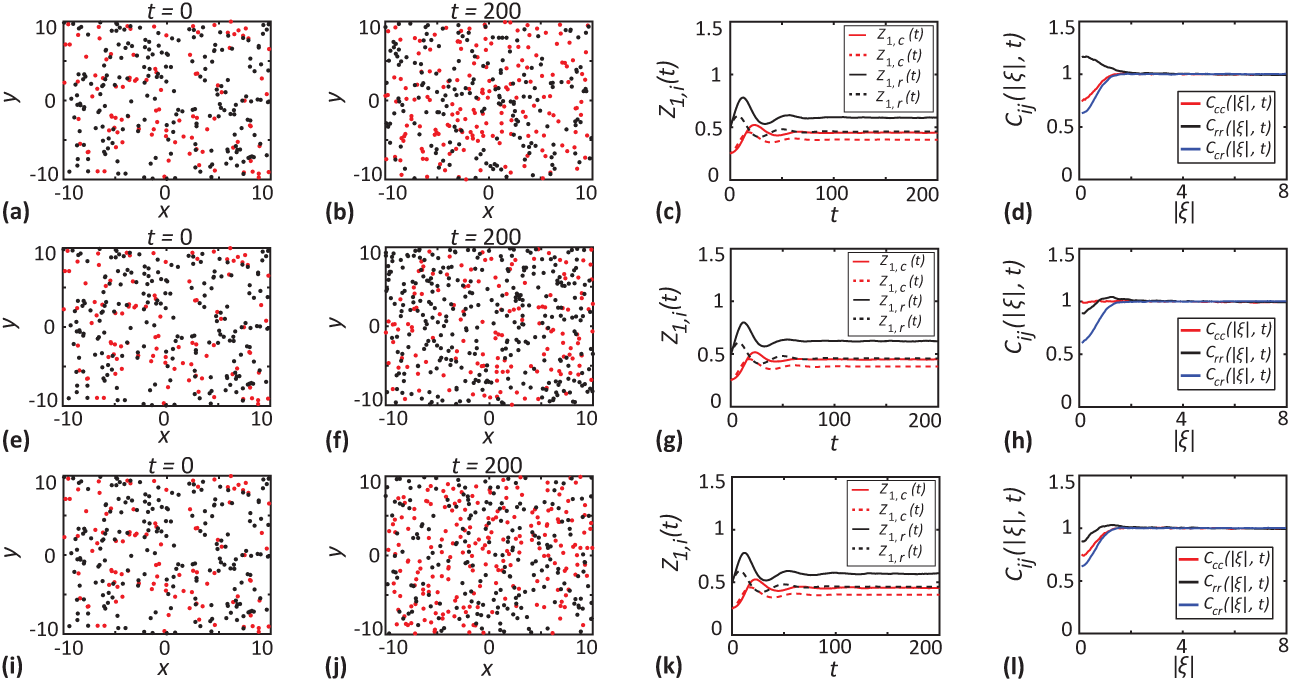
Short-range competition and short-range predation. Results in **a-d** correspond to short range competition of consumers (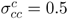and 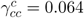) and short-range predation 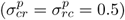. Results in **e-h** correspond to short-range competition of resources (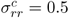 and 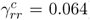) and short-range predation. Results in **i-l** correspond to short-range competition of both species (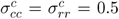 and 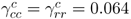) and short-range predation. **a, e, i** show the locations of consumers (red dots) and resources (black dots) at *t* = 0. **b, f, j** show the locations of consumers and resources at *t* = 200. **c, g, k** show the densities of consumers (red lines) and resources (black lines) as a function of time. Solid lines correspond to the averaged results from 1000 realisations of the IBM and dashed lines correspond to the solution of the mean-field model, Equations (6)-(7). **d, h, l** show the PCFs computed at *t* = 200 as a function of separation distance. All other parameters are same as in Table 1.

Combined effects of short-range competition of resources and short-range predation are shown in Figure 7(d)–(f). These conditions lead to interspecies segregation due to short-range predation. In addition, we see less pronounced intraspecies segregation developing among resources due to the competition between resources. In Figure 5, we saw that the effect of short-range predation, when considered in isolation, leads to a small scale clustering of resources. Here we see that the additional effects of predation counteract the impact of competition among resources to form a less pronounced intraspecies segregation of resources. Finally, we consider the combined effect of the short-range competition of both consumers and resources and short-range predation in Figure 7(g)–(i). We find competition-induced intraspecies segregation in this case, and this is confirmed by *C*_*cc*_(|***ξ***|, *t*) < 1 and *C*_*rr*_(|***ξ***|, *t*) < 1, respectively. We also observe interspecies segregation between consumers and resources confirmed by *C*_*cr*_(|***ξ***|, *t*) < 1 due to the short-range predation.

### 3.6 Spatial structure drives qualitative departure from the mean-field model

In this final set of simulations, we present a dramatic case where three spatial structure forming mechanisms act in unison to produce results that are fundamentally at odds with the predictions of the mean-field model. These results highlight the danger in neglecting spatial structure since the mean-field model leads to dramatically misleading predictions in this case.

The density dynamics of consumers and resources from the IBM simulation in Figure 8(a) show the eventual extinction of consumers while the resource species continues to grow to a steady density. In stark contrast, the solution of the mean-field model predicts the long-time coexistence of consumers and resources. These results demonstrate that the mean-field model and IBM can predict two qualitatively different outcomes when there is a strong spatial structure. In Figure 8(b)–(g), we show the evolution of the spatial structure of the community by presenting snapshots from the IBM as well as the PCFs at various points in time. We see intraspecies clustering among both consumers and resources, confirmed by *C*_*cc*_(|***ξ***|, *t*) > 1 and *C*_*rr*_(|***ξ***|, *t*) > 1, and interspecies segregation, confirmed by *C*_*cr*_(|***ξ***|, *t*) < 1 at earlier times before reaching the long-time limit. Strong clustering at these intermediate times is due to the short-range dispersal of both consumers and resources 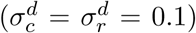. Competition tends to reduce the extent of clustering. But, dispersal dominates competition since we use very small dispersal range compared to the competition range 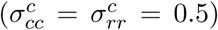. Short-range predation separate consumers and resources from each other by the consumption of close-lying resources by consumers. This results in interspecies segregation. As time progresses, these effects result in consumers becoming isolated from resources and eventually go extinct.

**Figure 8:**
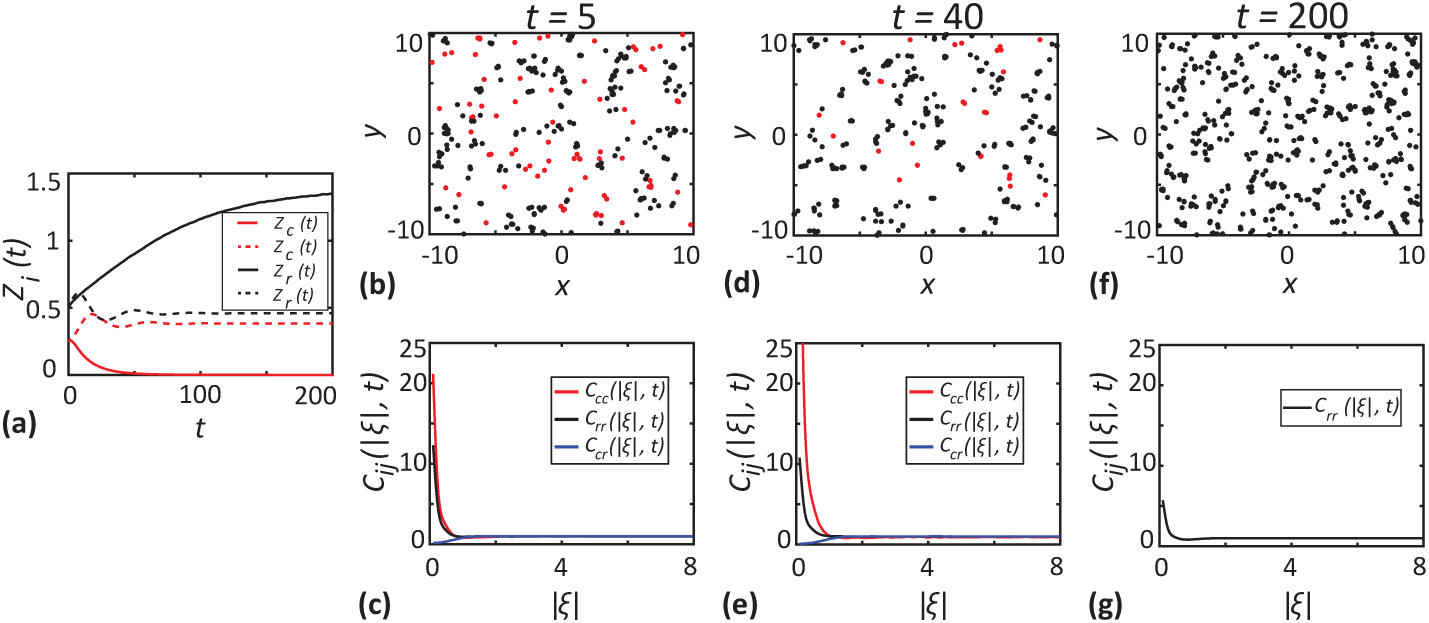
Short-range competition, short-range predation and short-range dispersal. **a** shows the densities of consumers (red lines) and resources (black lines) as a function of time. Solid lines correspond to the averaged results from 1000 realisations of the IBM and dashed lines correspond to the solution of the mean-field model, Equations (6)-(7). **b-c** show the locations of consumers (red dots) and resources (black dots) at *t* = 5 and PCFs computed at *t* = 5 as a function of separation distance. **d-e** show the locations of consumers and resources at *t* = 40 and PCFs computed at *t* = 40 as a function of separation distance. **f-g** show the locations of consumers and resources at *t* = 200 and PCFs computed at *t* = 200 as a function of separation distance. Parameter values are 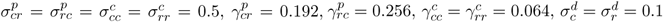. All other parameters are same as in Table 1.

## 4 Conclusion

In this work, we present a relatively simple IBM describing the dynamics of a community of consumers and resources where we pay particular attention to the effects of short-range interactions and small-scale spatial structure. The IBM simulation results are compared with solutions of a mean-field model constructed by invoking the mean-field approximation. This analysis reveals various situations where the IBM suggest qualitatively different outcomes compared to the solution of the mean-field model. These observations highlight the importance of considering the role of spatial structure in community dynamics. Mean-field models are routinely used to explore various characteristics of population dynamics and predator-prey type interactions. These include predicting whether certain subpopulations go extinct or whether all subpopulation survive and co-exist (Kuperman et al. 2019). Our results suggest that the predictions of the mean-field models in these scenarios could be inaccurate if the communities in question exhibit significant spatial structure.

Many studies consider the comparison of IBMs and their mean-field approximations to explore scenarios where mean-field assumption breaks down and predicts conflicting outcomes due to short-range interactions and spatial structure in a consumer resource community (Wilson 1998; Cantrell and Cosner 2004). While our present work explores cases where the mean-field model fails to predict coexistence and extinction accurately, similar studies in the literature consider other interesting scenarios where IBMs can exhibit stable dynamics when mean-field models predict oscillatory dynamics (Hosseini 2006; Brigatti et al. 2009). In these IBMs, limited motility and short-range interactions dampen the magnitude of population oscillations and stabilise the consumer-resource density dynamics (Cuddington and Yodzis 2000; Hosseini 2003). Another remarkable difference between our model and these previous modelling frameworks is here we use PCFs to identify the spatial configuration of the community. We base our inference about any observed changes in the density dynamics on findings revealed by PCFs about the spatial arrangement of the community. Additionally, we use a more realistic lattice-free IBM compared to the lattice-based models considered in these previous studies, which restrict the freedom of movement along fixed lattice directions.

Our modelling framework can be extended by incorporating various additional forms of neighbour-dependent interactions that are not considered here. An interesting extension would be to consider the effects of neighbour-dependent directional bias on the movement of consumers and resources. For example, in this work we make the simple assumption that the motility rate and direction of movement are independent of the local density. However, some recent IBMs introduce attractive or repulsive interactions where the motility rate and direction is affected by the local density, and these interactions can also drive significant differences in the macroscale spatial structure (Browning et al. 2018, 2020; Surendran et al. 2019; Binny et al. 2020). Incorporating these types of directional bias to the movement of consumers and resources may further influence the community dynamics beyond the effects explored in the present study. Another important neighbour-dependent effect in the context of population dynamics in ecology and cell biology is the incorporation of an Allee effect where proliferation and death rates are incorporated into the IBM so that there can be a net negative growth rate at low density and a net positive growth rate at higher densities (Stephens et al. 1999; Johnston et al. 2017; Fadai et al. 2019). Understanding how these kinds of interactions that lead to Allee effects would influence the formation of spatial structure is an open question that could be analysed by extending the modelling framework that we have presented here.

## Data availability

Matlab code used to generate results are available on Github at https://github.com/Anudeep-Surendran/Surendran2020

## acknowledgements

This work is supported by the Australian Research Council (DP170100474). MJP is partly supported by Te Pūnaha Matatini, a New Zealand Centre of Research Excellence. We thank the Associate Editor and anonymous referees for their helpful suggestions.

## References

[1] Abrams PA (2000) The evolution of predator-prey interactions: theory and evidence. Annual Review of Ecology and Systematics 31: 79–105.

[2] Abrams PA, Ginzburg LR (2000) The nature of predation: prey dependent, ratio dependent or neither? Trends in Ecology and Evolution 15: 337–341.

[3] Agnew DJG, Green JEF, Brown TM, Simpson MJ, Binder BJ (2014) Distinguishing between mechanisms of cell aggregation using pair-correlation functions. Journal of Theoretical Biology 352: 16–23.

[4] Akira S, Uematsu S, Takeuchi O (2006) Pathogen recognition and innate immunity. Cell 124: 783–801.

[5] Baker RE, Simpson MJ (2010) Correcting mean-field approximations for birth-death-movement processes. Physical Review E. 82: 041905.

[6] Barraquand F, Murrell DJ (2012) Intense or spatially heterogeneous predation can select against prey dispersal. PLoS One 7: e28924

[7] Binder BJ, Simpson MJ (2015) Spectral analysis of pair-correlation bandwidth: application to cell biooogy images. Royal Society Open Science 2: 140494.

[8] Binny RN, Haridas P, James A, Law R, Simpson MJ, Plank MJ (2016a) Spatial structure arising from neighbour-dependent bias in collective cell movement. PeerJ 4: e1689.

[9] Binny RN, James A, Plank MJ (2016b) Collective cell behaviour with neighbour-dependent proliferation, death and directional bias. Bulletin of Mathematical Biology 78: 2277–2301.

[10] Binny RN, Law R, Plank MJ (2020) Living in groups: Spatial-moment dynamics with neighbour-biased movements. Ecological Modelling 415: 108825.

[11] Binny RN, Plank MJ, James A (2015) Spatial moment dynamics for collective cell movement incorporating a neighbour-dependent directional bias. Journal of the Royal Society Interface 12: 20150228.

[12] Bolker BM, Pacala SW (1999) Spatial moment equations for plant competition: understanding spatial strategies and the advantages of short dispersal. The American Naturalist 153: 575–602.

[13] Brigatti E, Oliva M, Nunez-Lopez M, Oliveros-Ramos R, Benavides J (2009) Pattern formation in a predator-prey system characterized by a spatial scale of interaction. Europhysics Letters 88: 68002.

[14] Britton N (2003) Essential mathematical biology. Springer, London.

[15] Browning AP, McCue SW, Binny RN, Plank MJ, Shah ET, Simpson MJ (2018) Inferring parameters for a lattice-free model of cell migration and proliferation using experimental data. Journal of Theoretical Biology 437: 251–260.

[16] Browning AP, Jin W, Plank MJ, Simpson MJ (2020) Identifying density-dependent interactions in collective cell behaviour. bioRxiv. Accessed March 2020.

[17] Cantrell RS, Cosner C (2004) Deriving reaction–diffusion models in ecology from interacting particle systems. Journal of Mathematical Biology 48: 187–217.

[18] Cuddington KM, Yodzis P (2000) Diffusion-limited predator-prey dynamics in euclidean environments: An allometric individual-based model. Theoretical Population Biology 58: 259–278.

[19] Dini S, Binder BJ, Green JEF (2018) Understanding interactions between populations: individual based modelling and quantification using pair correlation functions. Journal of Theoretical Biology 439: 50–64.

[20] Dobramysl U, Tauber UC (2013) Environmental versus demographic variability in two-species predator-prey models. Physical Review Letters 110: 048105.

[21] Edelstein-Keshet L (2005) Mathematical models in biology (classics in applied mathematics). Society for Industrial and Applied Mathematics, New York.

[22] Fadai NT, Johnston ST, Simpson MJ (2019) Unpacking the Allee effect: determining individual-level mechanisms that drive population dynamics. bioRxiv. Accessed December 2019.

[23] Galetti M, Moleon M, Jordano P, Pires MM, Guimaraes PR Jr, Pape T, Nichols E, Hansen D, Olesen JM, Munk M, de Mattos JS, Schweiger AH, Owen-Smith N, Johnson CN, Marquis RJ, Svenning JC (2018) Ecological and evolutionary legacy of megafauna extinctions. Biological Reviews 93: 845–862.

[24] Gerum R, Richter S, Fabry B, Bohec CL, Bonadonna F, Nesterova A, Zitterbart DP (2018) Structural organisation and dynamics in king penguin colonies. Journal of Physics D: Applied Physics 51: 164004.

[25] Gillespie DT (1977) Exact stochastic simulation of coupled chemical reactions. The Journal of Physical Chemistry 81: 2340–2361.

[26] Grunbaum D (2012) The logic of ecological patchiness. Interface Focus 2: 150–155.

[27] Hillen T, Painter KJ (2009) A user’s guide to PDE models for chemotaxis. Journal of Mathematical Biology 58: 183–217.

[28] Hosseini PR (2003) How localized consumption stabilizes predator-prey systems with finite frequency of mixing. The American Naturalist 161: 567–585.

[29] Hosseini PR (2006) Pattern formation and individual-based models: The importance of understanding individual-based movement. Ecological Modelling 194: 357–371.

[30] Hunt VM, Brown JS (2018) Coexistence and displacement in consumer-resource systems with local and shared resources. Theoretical Ecology 11: 83–93.

[31] Jin W, McCue SW, Simpson MJ (2018) Extended logistic growth model for heterogeneous populations. Journal of Theoretical Biology 445: 51–61.

[32] Johnston ST, Baker RE, McElwain DLS, Simpson MJ (2017) Co-operation, competition and crowding: a discrete framework linking Allee kinetics, nonlinear diffusion, shocks and sharp-fronted travelling waves. Scientific Reports 7: 42134.

[33] Kuperman MN, Laguna MF, Abramson G, Monjeau JA (2019) Meta-population oscillations from satiation of predators. Physica A 527: 121288

[34] Law R, Dieckmann U (2000) A dynamical system for neighbourhoods in plant communities. Ecology 81: 2137–2148.

[35] Law R, Illian J, Burslem DFRP, Gratzer G, Gunatilleke CVS, Gunatilleke IAUN (2009) Ecological information from spatial patterns of plants: insights from point process theory. Journal of Ecology 97: 616–628.

[36] Law R, Murrell DJ, Dieckmann U (2003) Population growth in space and time: Spatial logistic equations. Ecology 84: 252–262.

[37] Markham DC, Simpson MJ, Maini PK, Gaffney EA, Baker RE (2013) Incorporating spatial correlations into multispecies mean-field models. Physical Review E 88: 052713.

[38] Mathworks (2019) Solve nonstiff differential equations — medium order method.

[39] Mobilia M, Georgiev IT, Tauber UC (2006) Fluctuations and correlations in lattice models for predator-prey interaction. Physical Review E 73: 040903(R).

[40] Mobilia M, Georgiev IT, Tauber UC (2007) Phase transitions and spatio-temporal fluctuations in stochastic lattice Lotka–Volterra models. Journal of Statistical Physics 128: 447–483.

[41] Murray JD (1989) Mathematical biology. Springer, New York.

[42] Murrell DJ (2005) Local spatial structure and predator-prey dynamics: counter-intuitive effects of prey enrichment. The American Naturalist 166: 354–367.

[43] Ovaskainen O, Finkelshtein D, Kutoviy O, Cornell S, Bolker B, Kondratiev Y (2014) A general mathematical framework for the analysis of spatiotemporal point processes. Theoretical Ecology 7: 101–113.

[44] Penczykowski RM, Laine A, Koskella B (2016) Understanding the ecology and evolution of host–parasite interactions across scales. Evolutionary Applications 9: 37–52.

[45] Plank MJ, Law R (2015) Spatial point processes and moment dynamics in the life sciences: A parsimonious derivation and some extensions. Bulletin of Mathematical Biology 77: 586–613.

[46] Plank MJ, Simpson MJ, Binny RN (2019) Small-scale spatial structure influences large-scale invasion rates. Theoretical Ecology https://doi.org/10.1007/s12080-020-00450-1.

[47] Rincon DF, Canas LA, Hoy CW (2017) Modeling changes in predator functional response to prey across spatial scales. Theoretical Ecology 10: 403–415.

[48] Santora JA, Reiss CS, Loeb VJ, Veit RR (2010) Spatial association between hotspots of baleen whales and demographic patterns of Antarctic krill Euphausia superba suggests size-dependent predation. Marine Ecology Progress Series 405: 255–269.

[49] Soehnlein O, Steffens S, Hidalgo A, Weber C (2017) Neutrophils as protagonists and targets in chronic inflammation. Nature Reviews Immunology 17: 248–261.

[50] Stephens PA, Sutherland WJ, Freckleton RP (1999) What is the Allee effect? Oikos 87: 185–190.

[51] Surendran A, Plank MJ, Simpson MJ (2018) Spatial moment description of birth-death-movement processes incorporating the effects of crowding and obstacles. Bulletin of Mathematical Biology 80: 2828–2855.

[52] Surendran A, Plank MJ, Simpson MJ (2019) Spatial structure arising from chase-escape interactions with crowding. Scientific Reports 9: 14988.

[53] Tobin P, Bjornstad ON (2003) Spatial dynamics and cross-correlation in a transient predator-prey system. Journal of Animal Ecology 72: 460–467.

[54] Treloar KK, Simpson MJ, Binder BJ, McElwain DLS, Baker RE (2015) Assessing the role of spatial correlations during collective cell spreading. Scientific Reports 4: 5713.

[55] Vijay K (2018) Toll-like receptors in immunity and inflammatory diseases: Past, present, and future. International Immunopharmacology 59: 391–412.

[56] Wang H, Nagy JD, Gilg O, Kuang Y (2009) The roles of predator maturation delay and functional response in determining the periodicity of predator–prey cycles. Mathematical Biosciences 221: 1–10.

[57] Wilson WG (1998) Resolving discrepancies between deterministic population models and individual-based simulations. The American Naturalist 151: 116–134.

